# Proliferation of SH-SY5Y neuroblastoma cells on confined spaces

**DOI:** 10.1101/2024.02.05.578902

**Authors:** Ewelina Kalwarczyk, Agnieszka Lukasiak, Damian Woznica, Weronika Switlik, Julia Anchimowicz, Piotr Zielonka, Slawomir Jakiela

## Abstract

**Background:** Microfluidics offers precise drug delivery and continuous monitoring of cell functions, which is crucial for studying the effects of toxins and drugs. Ensuring proper cell growth in these space-constrained systems is essential for obtaining consistent results comparable to standard Petri dishes.

**New method:** We investigated the proliferation of SH-SY5Y cells on circular polycarbonate chambers with varying surface areas. SH-SY5Y cells were chosen for their relevance in neurodegenerative disease research.

**Results:** Our study demonstrates a correlation between the chamber surface area and SH-SY5Y cell growth rates. Cells cultured in chambers larger than 10 mm in diameter exhibited growth comparable to standard 60-mm dishes. In contrast, smaller chambers significantly impeded growth, even at identical seeding densities. Similar patterns were observed for HeLaGFP cells, while 16HBE14*σ* cells proliferated efficiently regardless of chamber size. Additionally, SH-SY5Y cells were studied in a 12-mm diameter sealed chamber to assess growth under restricted gas exchange conditions.

**Comparison with existing methods:** Our findings underscore the limitations of small chamber sizes in microfluidic systems for SH-SY5Y cells, an issue not typically addressed by conventional methods.

**Conclusions:** SH-SY5Y cell growth is highly sensitive to spatial constraints, with markedly reduced proliferation in chambers smaller than 10 mm. This highlights the need to carefully consider chamber size in microfluidic experiments to achieve cell growth rates comparable to standard culture dishes. The study also shows that while SH-SY5Y and HeLaGFP cells are affected by chamber size, 16HBE14*σ* cells are not. These insights are vital for designing effective microfluidic systems for bioengineering research.

## Introduction

Traditional microscopic studies in cell biology and bioengineering use fixed biological preparations, which are the gold standard for assessing the impact of compounds on cell function and viability (1). However, this approach inherently limits the ability to observe dynamic processes occurring in real time. Using microfluidic technology, we can easily combine microscopic inspection with precise drug delivery (2, 3). This breakthrough makes it possible to continuously monitor the influence of different toxins or drugs on cell function (4). To go even further, organ-on-a-chip systems function like real tissue systems and show how they react to disease (5, 6). The authors wanted to use microfluidic techniques to study SH-SY5Y cells, which have been chosen as a model for neurodegenerative diseases (7, 8). In basic research into neurodegenerative diseases, the choice of an appropriate model system is crucial (9, 10). Both human and animal cells are used, including primary and immortalised cell lines. The human immature neuroblastoma cell line SH-SY5Y, widely used in Parkinson’s and Alzheimer’s disease research (11, 12), can be modified to mimic dopaminergic (13, 14) or cholinergic (15, 16) states. These properties make it valuable for studying toxic protein aggregation (17, 18) and mitochondrial dysfunction (19, 20). However, in our initial tests, growing SH-SY5Y cells in small microfluidic chambers (polycarbonate, glass and polydimethylsiloxane cell substrates were tested) with dimensions as small as 0.5 mm slowed their growth rate more than 10-fold compared to standard dishes. We therefore carried out a series of calibration studies to identify the ideal chamber size to achieve cell proliferation similar to that observed in conventional culture dishes.

These studies give us very interesting information about how SH-SY5Y cells grow in carefully made polycarbonate chambers ranging in diameter from 3mm to 50mm. Importantly, we used these chambers without any additional surface treatment to enhance cell adhesion. A comparative analysis of cell growth in these chambers versus conventional 60 mm diameter Petri dishes (Corning) was performed. All images capturing the coverage of the test media during successive days of culture are available in the Zenodo repository (21). Real-time monitoring of cell growth was achieved using a phase-contrast microscope, and growth kinetics were methodically analysed by extensive examination of microscopic images. We also studied the respiration of SH-SY5Y cells using the Oroboros Oxygraph-2k device. The aim of this study was to explain how quickly a population of cells in a sealed chamber without gas exchange consumes oxygen in the culture medium. We studied the growth of SH-SY5Y cells over a one-week observation period in a hermetic polycarbonate chamber specifically designed for targeted studies in microfluidic systems. In addition, a video showing the growth of SH-SY5Y cell cultures, with individual culture frames recorded every 5 minutes, has been deposited in the Zenodo repository (22).

Research in cell biology and bioengineering is crucial, especially as we move towards the widespread use of small microfluidic systems (4, 23). However, these systems present significant challenges for cell function in confined spaces, a point highlighted in our study. The observed sensitivity of SH-SY5Y to spatial constraints highlights the need for careful consideration of chamber size in experiments aimed at modelling disease conditions, particularly in lab-on-a-chip and organ-on-a-chip systems. Understanding how these cells behave in a confined environment and providing specific solutions - in this case, the size of the experimental chambers - can help to design more effective microfluidic systems for studying neuronal behaviour and drug response. Our study does not only focus on SH-SY5y cells, but we also present results for HeLaGFP and 16HBE14*σ* cells to show that the problem presented is not unique to SH-SY5y cells.

## Materials and methods

### Materials

The following chemicals and solutions were provided by Sigma-Aldrich, Germany: Dulbecco’s modified Eagle’s medium (DMEM) - D6546, Eagle’s minimum essential medium (MEM) - M4655, heat inactivated fetal bovine serum (FBS) - F9665, L-glutamine solution, 200 mM - G7513, antibiotics solution containing 10 000 units of penicillin, 10 mg/mL streptomycin - P0781, MEM Non-Essential Amino Acid (NEAA) - M7145, Trypsin-EDTA solution - T3924, sodium dodecyl sulphate (SDS) - L3771, dichloromethene – 650463, Oligomycin - O4876, carbonyl cyanide 4-(trifluoromethoxy)phenylhydrazone (FCCP)- C2920, rotenone - R8875, and antimycin A - A8674. Ethyl alcohol, 96% was obtained from POCH, Poland (396420113), phosphate buffered saline (PBS) without calcium and magnesium was purchased from Lonza.

### Cells

Three cell lines were used to study cellular responses in a controlled laboratory environment. The first cell line, SH-SY5Y human neuroblastoma, was obtained from ECACC and cultured in DMEM with 10% FBS, 1% 200 mM L-glutamine, 1% antibiotic solution and 1% NEAA for optimal growth and viability.

In addition, the study included the Milipore 16HBE14*σ* human bronchial epithelial cell line, which was cultured in MEM with 10% FBS and 1% antibiotic solution to diversify cellular components and increase the breadth of the study.

A third cell line, HeLaGFP inducible GFP HeLa stable, purchased from GenTarget Inc., was grown in DMEM with 10% FBS, 1% 200 mM L-glutamine and 1% antibiotics solution. Cell growth was continuously monitored using an Olympus CKX53 phase-contrast microscope with a DLT camera (Delta Optical, DLT-Cam PRO 1080 HDMI USB) in a temperature-controlled chamber set at 37 °C according to experimental requirements.

### Chambers

In our investigation, cell cultures were grown in a variety of vessels, including commercially available cell culture dishes such as NUNC Easy Flasks (75 cm^2^) from Thermo Scientific. We also used Corning 60 mm Petri dishes. In a more tailored approach, our research involved the use of specially designed multi-chamber cultures (see Figure 1). These chambers were carefully and precisely fabricated by milling 10 mm and 5 mm thick polycarbonate plates (Makrolon, Bayer, Leverkusen, Germany) using a state-of-the-art CNC milling machine (MSG4025, Ergwind, Gdansk, Poland). To reduce evaporation of the medium and to mimic the conditions of a standard Petri dish for cell culture, these chambers were fitted with a polycarbonate lid.

**Fig. 1.**
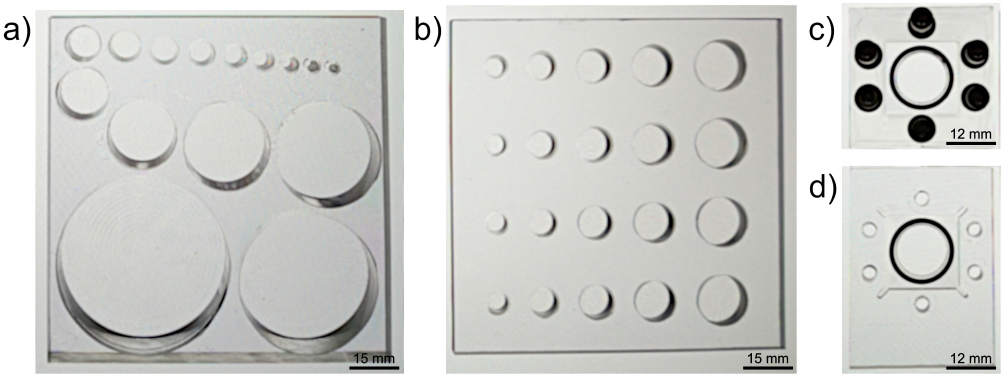
Photographs depicting a custom-designed multi-chamber culture dish, which has been carefully manufactured from polycarbonate using CNC milling, are presented here. The culture dish consists of the following components: a) A 15-well multichamber with varying chamber diameters ranging from 2.4 to 50 mm. The diameters include 2.4, 3, 4, 5, 6, 7, 8, 9, 10, 15, 20, 25, 30, 40 and 50 mm. b) A 20-well chamber with diameters of 6, 8, 10, 12 and 15 mm. Note that each diameter is replicated four times to facilitate comparative experiments. c) The top layer and d) the bottom layer of a hermetically sealed 12 mm diameter polycarbonate chamber are also shown. These chambers were sealed with an O-ring and held together with 6 M3 screws.

We decided to use polycarbonate as a superior choice for low-cost cell culture experiments due to its excellent biocompatibility(24). The biologically inert nature of polycarbonate ensures that cell viability and behaviour are unaffected, a critical factor in maintaining healthy cell cultures over long periods of time (25). The optical properties of polycarbonate make it an ideal substrate for cell culture, providing conditions similar to standard procedures using glass or polystyrene Petri dishes, which is advantageous for the highquality imaging essential for real-time microscopy monitoring of cell dynamics (26). In addition, the ease of fabrication and modification of polycarbonate allows the creation of customised multi-chamber culture dishes with different surface areas using techniques such as laser cutting (27), milling and thermal bonding (28).

### Statistical analysis

Quantitative experiments were performed in triplicate and data are presented as mean ± standard error of the mean. One-way analysis of variance (ANOVA) was used to compare different experimental groups. Post-hoc comparisons with the control group (typically a 60 mm Petri dish) were performed using the Dunnett’s test; therefore, ‘adj. p’ is applied in the description of the results, meaning ‘adjusted p-value’. Statistical analyses and graphs were generated using GraphPad Prism software.

## Experimental

### Fabrication of multi-chamber cell dishes

To improve the quality of the prepared culture dishes, their surfaces were smoothed using a stream of methylene chloride vapour. They were then carefully cleaned using a solution of 0.1M SDS, ethyl alcohol and deionised water. Finally, the dishes underwent a comprehensive sterilisation procedure, which included baking in an oven at 110 °C for 10 hours and exposure to UV light for 30 minutes in a cell culture hood.

### Growth of SH-SY5Y cells in a polycarbonate culture dish

In this study, a freshly prepared cell suspension, characterised by a known cell concentration per millilitre using Bio-Rad’s TC20 automated cell counter, was evenly distributed across all chambers of the culture dish. The volume of the cell suspension was carefully adjusted to maintain a consistent liquid height of approximately 2 mm in each chamber, ensuring a uniform cell surface density, measured in cells per square centimetre.

The prepared culture dishes were then placed in a cell culture incubator set at 37 °C with a 5% CO_2_ concentration. The growth of the culture was systematically observed in all the polycarbonate chambers, as well as in the other bottles and dishes mentioned, using a phase-contrast microscope. For each culture plate, five microscope images were taken - one at the centre of the plate and four at random locations. After image acquisition, the culture plate was returned to the incubator, allowing continuous and prolonged monitoring of cell growth. Microscopic image analysis allowed the average surface coverage of the culture plate to be determined over time. This analytical approach remained consistent throughout the majority of the experiments, unless explicitly stated otherwise.

The culture medium remained unchanged throughout the growth process, which typically lasted up to 7 days, unless specific changes were required for a particular experiment.

### Measurement of Oxygen Consumption Rate (OCR)

To elucidate the oxygen consumption rate (OCR) in SH-SY5Y cells, we used a Clark-type oxygen electrode according to the mitochondrial function assessment protocol described in a previously published methodology (29). SH-SY5Y cells were grown in NUNC Easy Flasks (75 cm^2^), each containing 12 ml of culture medium, to approximately 90% confluence (equivalent to approximately 10 million cells per flask). A suspension of SH-SY5Y cells in culture medium at concentrations of 1 million/ml and 2.5 million/ml was then added to the measuring chambers of the Oxygraph-2K system (Oroboros Instruments, Austria).

For the determination of the non-phosphorylating leak, 4 mg/ml oligomycin, an inhibitor of mitochondrial ATP synthase, was used to reflect the maximal coupled state of the mitochondria. To achieve maximum uncoupling of the electron transfer system (ETS), 1 mM FCCP (carbonyl cyanide 4-(trifluoromethoxy)phenylhydrazone) was used. Residual oxygen consumption was then determined by adding 2.5 mM rotenone and 0.5 mM antimycin A, inhibitors of complex I and III of the ETS, respectively.

The expected oxygen consumption time was estimated from the initial oxygen concentration values and the baseline OCR of the cells.

### Methodology of data collection

To characterise the developmental trajectory of cells within chambers of different dimensions, cell images taken on different days of culture were binarised. This was done either manually using the GNU Image Manipulation Program (GIMP) or semi-automatically using the Phantast phase-contrast microscopy segmentation toolbox in ImageJ. Phantast binarised images were then manually refined using the GIMP where necessary. Quantification of cell-covered areas was performed using ImageJ. The resulting values from each set of images were averaged, resulting in mean confluence measurements at different stages of the cell culture. An illustrative representation of SH-SY5Y cells under phase contrast microscopy is shown in Figure 2a, while the corresponding binarised representation is shown in Figure 2b.

**Fig. 2.**
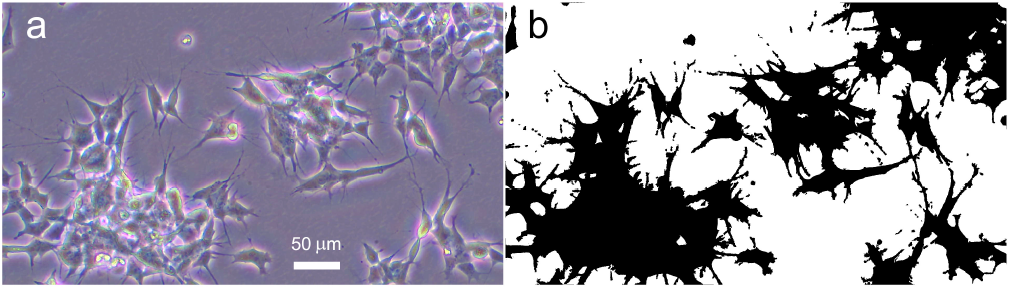
Phase-contrast microscope image of SH-SY5Y cells (a) together with the corresponding binarised image (b).

### Monitoring the growth of SH-SY5Y cells in a hermetically sealed polycarbonate chamber over a period of one week

This investigation uses biological microscopy techniques to provide real-time insights into the dynamic transformations and growth trajectories of SH-SY5Y cells within a carefully controlled microenvironment. The resulting images, taken at five-minute intervals, were processed and condensed into a comprehensive video using ImageJ, as detailed in the Zenodo repository (22). The hermetically sealed chamber design provides a precisely controlled environment, maintaining a constant temperature throughout the microscopic observation and eliminating the need to move the system. This design choice allows accurate monitoring and comprehensive documentation of cell behaviour, interactions and growth patterns over a predetermined one-week period.

The sealed chamber used, which is an integral part of the research, consists of two intricately interconnected polycarbonate components that are securely bolted together and sealed with O-rings - as shown in Figure 1c, d. Prior to assembly, all elements underwent a thorough cleaning process and were annealed at 110 °C for 10 hours. After extensive sterilisation under UV light for 30 minutes in a laminar clean bench, the individual components were assembled. A 0.3 ml chamber (12 mm diameter) was then filled with a cell suspension containing 1 x 10^5^ cells/ml. Prior to microscopy, the culture was pre-incubated at 37 °C, 5% CO_2_ for 24 hours. The chamber was securely positioned within the microscope chamber and a constant temperature of 37 °C was maintained without CO_2_ control. At this point, real-time monitoring of SH-SY5Y cell growth began.

## Results and discussion

### Exploring the effects of seeding concentration and chamber size on SH-SY5Y cell growth dynamics

Investigating the effect of seeding concentration on SH-SY5Y cell growth dynamics provides valuable insights into cell proliferation kinetics. This study examines the growth of SH-SY5Y cells cultured in 60 mm Petri dishes and PC chambers of different diameters (6, 8, 10, 12 and 15 mm) at different seeding concentrations.

In the Petri dish experiments, cells were seeded at three different concentrations (1 *×* 10^4^, 5 *×* 10^4^ and 1 *×* 10^5^ cells/ml), with four separate culture plates for each concentration to ensure robustness and reliability. Microscopic images were analysed to determine average surface coverage over time, revealing a clear relationship between seeding concentration and cell proliferation rates (Figure 1a). Higher seeding concentrations resulted in faster growth rates, with the highest concentration reaching full confluence in 7 days, whereas the lowest concentration took approximately 2 weeks.

This direct correlation between seeding concentration and proliferation rate is consistent with existing literature suggesting that higher seeding concentrations enhance cell growth due to increased cell-cell interactions and resource availability (30–33).

The study also investigated the effect of seeding concentration in chambers of different diameters (see Figure 1b) using concentrations of 5 *×* 10^4^, 1 *×* 10^5^ and 5.5 *×* 10^5^ cells/ml. The microscopic images were taken at intervals of 1, 2, 3 and 7 days. Confluence analysis of SH-SY5Y (see Figure 3b-d) cells revealed significant observations in cell growth rates as a function of cell size, despite the same initial concentration. In smaller chambers (6 mm), cell aggregation occurred after an initial healthy growth phase, regardless of seeding concentration. In medium-sized chambers (8-10 mm), higher seeding concentrations led to rapid growth followed by aggregation, although aggregation also occurred at lower seeding concentrations. For larger chambers (greater than 10 mm), a seeding concentration of 1 *×* 10^5^ cells/ml was optimal for maintaining long-term observation of cell growth without aggregation.

**Fig. 3.**
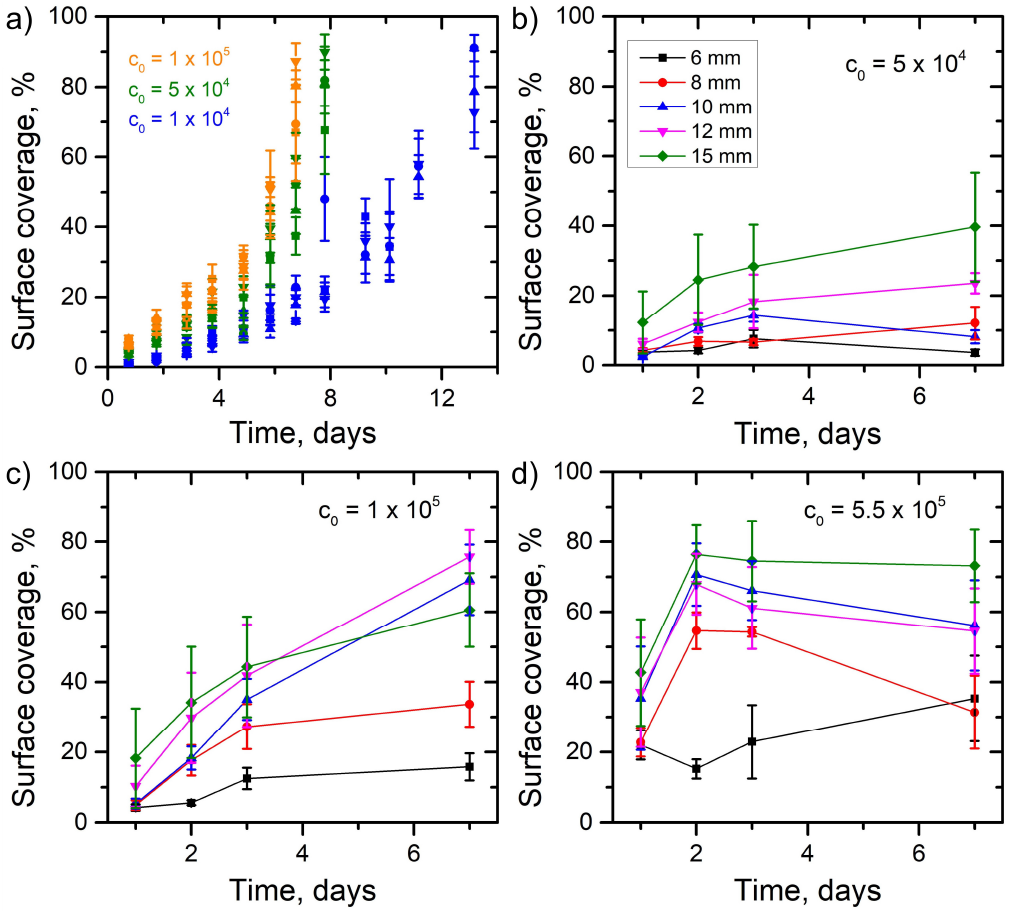
(a) The growth curve shows the proliferation of SH-SY5Y cells in a 60 mm culture plate, initially seeded at three different concentrations. (b) - (d): Growth curves of SH-SY5Y cells at different chamber diameters at different seeding concentrations. The colour-coded chamber diameters in panel (b) correspond to those in panels (c) and (d).

In conclusion, this comprehensive study provides powerful insights into the effects of seeding concentration and spatial constraints on the growth dynamics of SH-SY5Y cells. The results highlight the importance of optimising seeding density and chamber size to achieve desired growth outcomes, which is critical for improving cell culture practices and establishing consistent conditions for future experiments. The observed effect of chamber size on cell growth rates measured in only five different chamber diameters (see Figure 1b) prompted us to analyse the effect of chamber size on SH-SY5Y cell proliferation in detail.

### Investigation of the effect of available surface area on the growth of SH-SY5Y cells

To investigate the effect of available surface area on the growth dynamics of SH-SY5Y cells, we designed and fabricated multi-chamber polycarbonate culture dishes. These dishes had different surface areas ranging from 0.07 cm^2^ (in circular chambers of 3 mm diameter) to 19.63 cm^2^ (in circular chambers of 50 mm diameter), see Figure 1a. The preparation of these chambers, as described in the experimental section, ensured experimental integrity.

An accurate calculation of the cell suspension volume was made to achieve a uniform cell suspension height of 2 mm in each chamber, thereby establishing an equivalent average surface density of cells (e.g. for 1 *×* 10^4^ cells/cm^2^ the concentration was 5 *×* 10^4^ cells/ml). After seeding, the dish was securely covered with a polycarbonate lid and incubated in a controlled cell incubator (37 degrees C / 5% CO_2_). Using a phase-contrast microscope, we regularly assessed cell growth in the chambers every 1-2 days. The images captured showed dynamic changes in mean cell confluence over time.

Figures 4a-b show an interesting relationship between the growth curve of SH-SY5Y cells and the area used for cell growth. In smaller chambers, with a diameter of less than approximately 10 mm, the area coverage shows a proportional scaling with chamber size (see Figures 4a-d). The level of statistical significance for confluence comparisons of chambers smaller than 10 mm in diameter with a 60 mm Petri dish increases with time during cell culture (see Figures 4c-d). This indicates a much lower growth rate of SH-SY5Y cells for small chambers. It can be concluded that the smaller the chamber, the lower the growth rate for the same initial seeding concentration. There is no statistical significance in the area coverage when comparing chambers larger than 10 mm in diameter. For the chambers larger than 10 mm in diameter, area coverage appears to be independent of chamber size - lack of statistical significance in area coverage comparisons between chambers larger than 10 mm in diameter and a 60 mm Petri dish (see Figures 4c-d). In particular, the mean confluence remains consistently stable across all chambers larger than 10 mm in diameter.

**Fig. 4.**
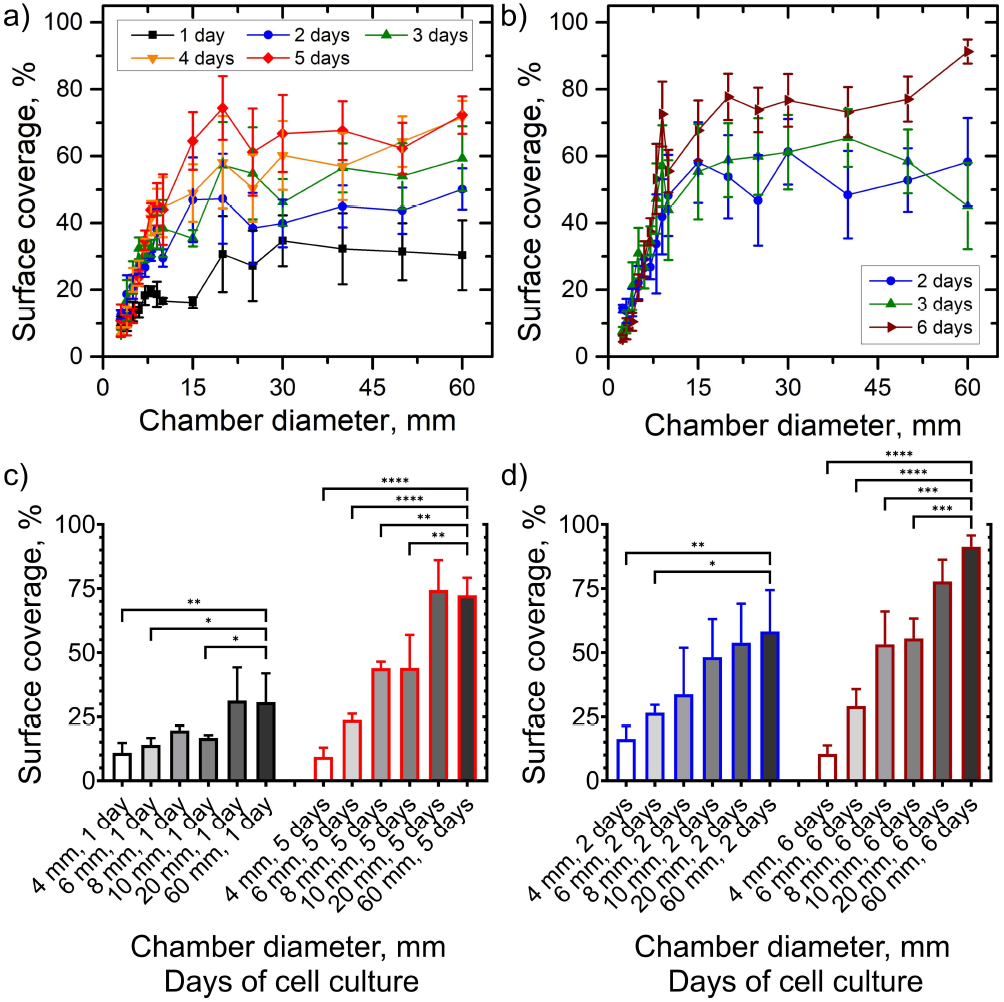
Variations in surface coverage within cell chambers were observed as a function of chamber diameter for SH-SY5Y cells. The specific conditions, denoted by passage number and initial cell concentration, were as follows: a) SH-SY5Y p10, with an initial concentration of 1 × 10^5^ cells/ml; b) SH-SY5Y p15, with an initial concentration of 5 × 10^4^ cells/ml. Guidelines have been included to aid visual interpretation. c-d) Statistics showing significance levels and their change for comparisons of area coverage for different size culture chambers with cell culture on a 60 mm Petri dish. For readability, only certain chamber sizes were selected. c) Concentration of 1 × 10^5^ cells/ml d) Concentration of 5 × 10^4^ cells/ml. Post hoc test: \**adj*.*p <* 0.05, * * *adj*.*p <* 0.01, * * \**adj*.*p <* 0.001, * * * * *adj*.*p <* 0.0001.

The above conclusions hold true for both initial cell seeding levels studied: 5 *×* 10^4^ cells/ml and 1 *×* 10^5^ cells/ml.

### The growth of HeLaGFP and 16HBE14*σ* cells in poly-carbonate multichamber

The growth of SH-SY5Y cells and their limited ability to adhere to surfaces compared to other cell types has been extensively studied by others. Numerous factors, including substrate stiffness, integrin-mediated signalling, growth factors and cell-cell interactions, have been identified as crucial in regulating the adhesive properties of SH-SY5Y cells (34, 35).

Motivated by these findings, we sought to compare the growth patterns of the more adherent 16HBE14*σ* and HeLaGFP cells within polycarbonate chambers (Figure 1a). The cell seeding process involved introducing cells from suspension at a concentration of 1 *×* 10^5^ cells/ml. Our study involved daily, detailed microscopic examinations to observe how the cells grew. The resulting growth curves, shown in Figure 5a-b, provide a comprehensive representation of the temporal evolution of cell proliferation within this experimental framework.

**Fig. 5.**
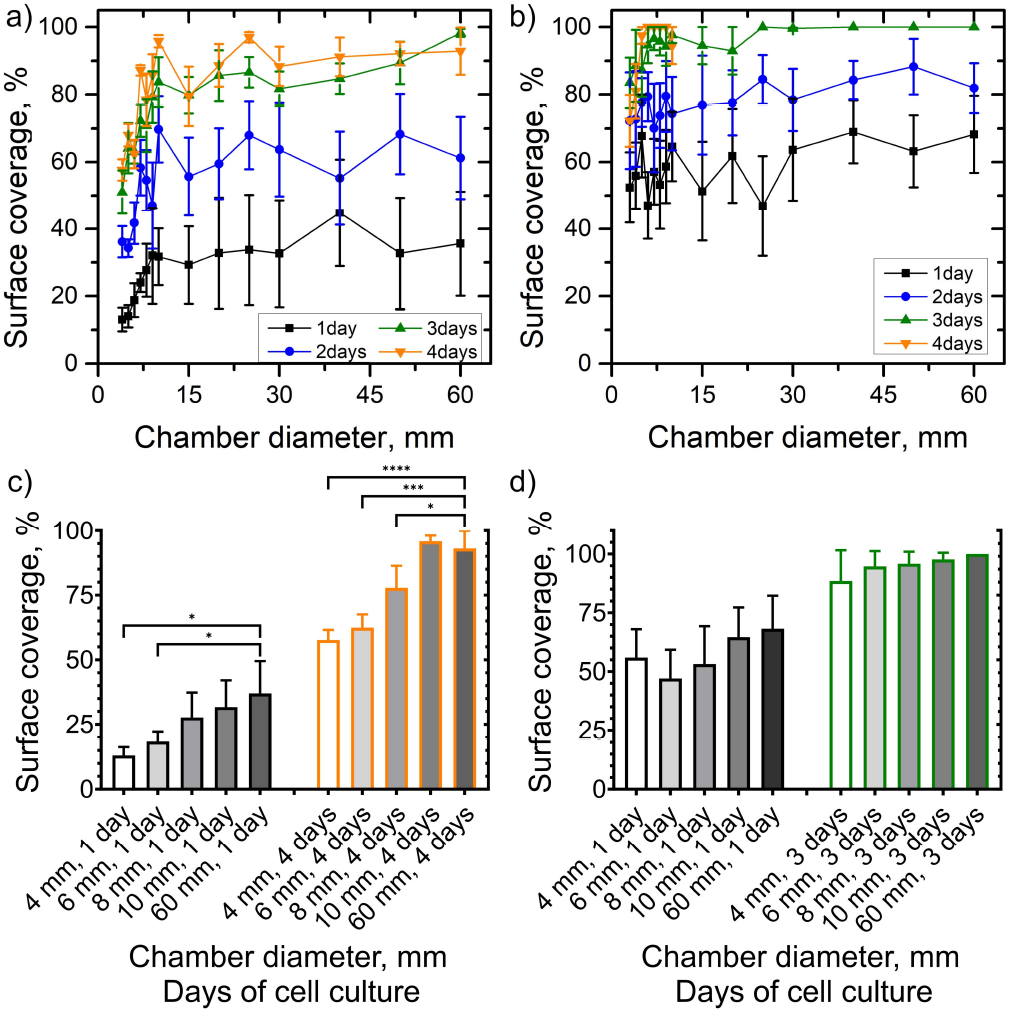
Temporal evolution of cell growth: tracking of the expansion of HeLaGFP (a) and 16HBE14*σ* (b) cell populations over time, plotted as surface coverage relative to chamber diameter. Cells were initially seeded at a concentration of 1 × 10^5^ cells/ml. c-d) Statistics showing significance levels and their change for comparisons of area coverage for different size culture chambers with cell culture on a 60 mm Petri dish. For readability, only certain chamber sizes were selected. c) HeLaGFP d) 16HBE14*σ*. Post hoc test: \**adj*.*p <* 0.05, * * *adj*.*p <* 0.01, * * \**adj*.*p <* 0.001, * * * * *adj*.*p <* 0.0001

Our study showed that the growth of HeLaGFP cells, like that of SH-SY5Y cells, is influenced by cell size. Chambers smaller than 10 mm had lower levels of HeLaGFP cell confluence compared to larger diameter chambers or cultures in 60 mm Petri dishes. These differences increased over time - the significance level of the coverage area for small chambers compared to the coverage area of a 60 mm Petri dish increased over time (see Figure 5c).

In contrast, 16HBE14*σ* cells showed no clear dependence of growth on chamber diameter - no statistical significance in confluence comparisons between chambers or when compared to cultures grown in a 60 mm Petri dish (see Figure 5d). Notably, smaller chambers (diameter < 6 mm) resulted in reduced coverage after 4 days compared to day 3 of culture for both cell lines. Nevertheless, both cell lines showed rapid growth compared to SH-SY5Y cells, reaching 90% confluence after 4 days. It can also be seen that 16HBE14*σ* cells showed faster growth in polycarbonate chambers compared to HeLaGFP cells.

### The SH-SY5Y cells show low mitochondrial function expressed as OCR

Mitochondrial functionality in SH-SY5Y cells is characterised by a low oxygen consumption rate (OCR), as shown in Figure 6. The reduced respiratory activity in SH-SY5Y cells is likely due to their unique neuronal characteristics, increased sensitivity to oxidative stress and possible mitochondrial problems. This is evidenced by lower levels of ATP, mitochondrial membrane potential (MMP) and citrate synthase activity(36, 37).

**Fig. 6.**
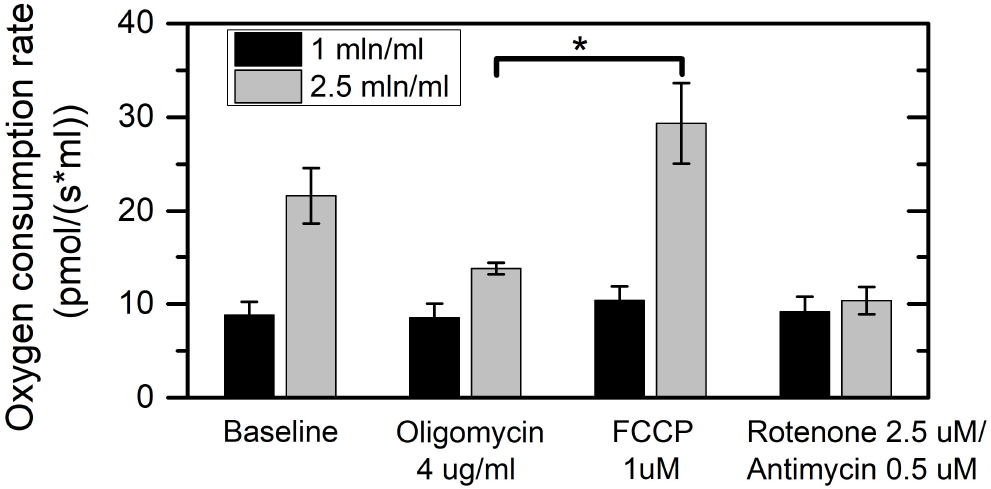
Oxygen consumption rate in SH-SY5Y cells: The depicted bars illustrate the OCR for two cell concentrations — 1 × 10^6^ cells/ml (black bars) and 2.5 × 10^6^ cells/ml (grey bars). Results are presented as mean ± standard error of the mean (SEM), with a sample size of *n* = 4 for the former and *n* = 3 for the latter. Statistical significance, denoted by (*), was determined at p < 0.05 using one-way analysis of variance (ANOVA).

In this study, we sought to elucidate the oxygen consumption dynamics of SH-SY5Y cells during the process of cellular respiration. The baseline OCR for a concentration of 1 million SH-SY5Y cells per millilitre was determined to be approximately 8.8 pmol/(s*ml). Following the introduction of oligomycin, an ATP synthase inhibitor, we restricted the flow of protons back into the mitochondrial matrix. This resulted in a significant reduction in cellular oxygen consumption, with OCR falling to 8.5 pmol/(s*ml). Following administration of FCCP, a compound known to uncouple the mitochondrial electron transport system (ETS), cellular respiration peaked at 10.4 pmol/(s*ml). Statistical analysis revealed no significant difference between the effects induced by FCCP and the baseline OCR or the OCR observed during ETS with inhibited ATP synthase (after oligomycin administration). In particular, using a 2.5-fold higher cell concentration, the OCR following FCCP administration exceeded that following oligomycin administration by more than twofold. Crucially, the baseline OCR showed a 2.5-fold escalation with increasing cell concentration, confirming a linear correlation between OCR and cell concentration. In the final phase of the cellular respiration study, we evaluated the inhibition of complexes I and III in the ETS using rotenone and antimycin A, respectively. Used in tandem, these compounds effectively impeded the flow of electrons throughout the electron transport chain, facilitating the measurement of extra-mitochondrial oxygen consumption.

The expected duration of oxygen consumption for different concentrations of SH-SY5Y cells is shown in table 1. Specifically, the approximate time for oxygen consumption is 29 days for a cell concentration of 10^4^ cells/ml and 2.9 days for a concentration of 10^6^ cells/ml. These results are consistent with the baseline Oxygen Consumption Rate (OCR) exhibited by the cells as shown in Figure 6. It is important to note that the estimated oxygen consumption time was derived from a linear relationship and did not control for cell division. To improve the accuracy of our understanding of oxygen consumption, it is important to include the rate of cell division in the calculation of the corrected OCR. In addition, the diffusion of oxygen within the polycarbonate system should be carefully considered to ensure a more accurate representation of the overall oxygen dynamics in the experimental setup.

**Table 1.** Expected duration of oxygen utilisation by SH-SY5Y cells at different concentrations in the culture medium, assuming no air exchange. The table shows results averaged to two significant digits. Abbreviations: O_2_ conc. - initial oxygen concentration in the culture medium; OCR - oxygen consumption rate; OCT - oxygen consumption time.

| | $10^4$ cells/ml | $10^6$ cells/ml |
| --- | --- | --- |
| $O_2$ conc. (pmol/ml) | 220000 | 220000 |
| OCR (pmol/s*ml) | 0.088 | 8.8 |
| OCT (s) | 2500000 | 25000 |
| OCT (min.) | 42000 | 420 |
| OCT (h) | 700 | 7 |
| OCT (days) | 29 | 0.29 |

### Long, real-time microscopic observation of the growth of SH-SY5Y cells in sealed chamber

In the early stages of culturing SH-SY5Y cells, the authors conducted preliminary tests using small microfluidic chambers with a diameter of 0.5 mm. Unfortunately, there was no observable cell growth in these chambers. Subsequent studies showed that SH-SY5Y cells showed a marked preference for chambers larger than 10 mm in diameter. Based on these findings, a carefully designed sealed chamber with a diameter of 12 mm and a volume of 0.3 ml was constructed - see Figure 1c, d. Continuous microscopic monitoring of SH-SY5Y cell growth dynamics within this chamber was then performed.

The aim of this study was to precisely follow the real-time microscopic development of the cells without external gas exchange. The chamber volume of 0.3 ml was chosen based on the cellular respiration results and the requirement for observations over at least one week. Consequently, the sealed chamber has an initial cell surface density of 200 cells/mm^2^. Figure 7a shows how the surface coverage in the sealed chamber changes over time. Comparing a sealed 12 mm diameter chamber with a standard culture chamber, the area covered by cells was similar on days 2, 3 and 7 of the culture period, as shown in Figure 3c. This suggests that long-term cell culture without air exchange through the medium is comparable to conventional cell culture.

**Fig. 7.**
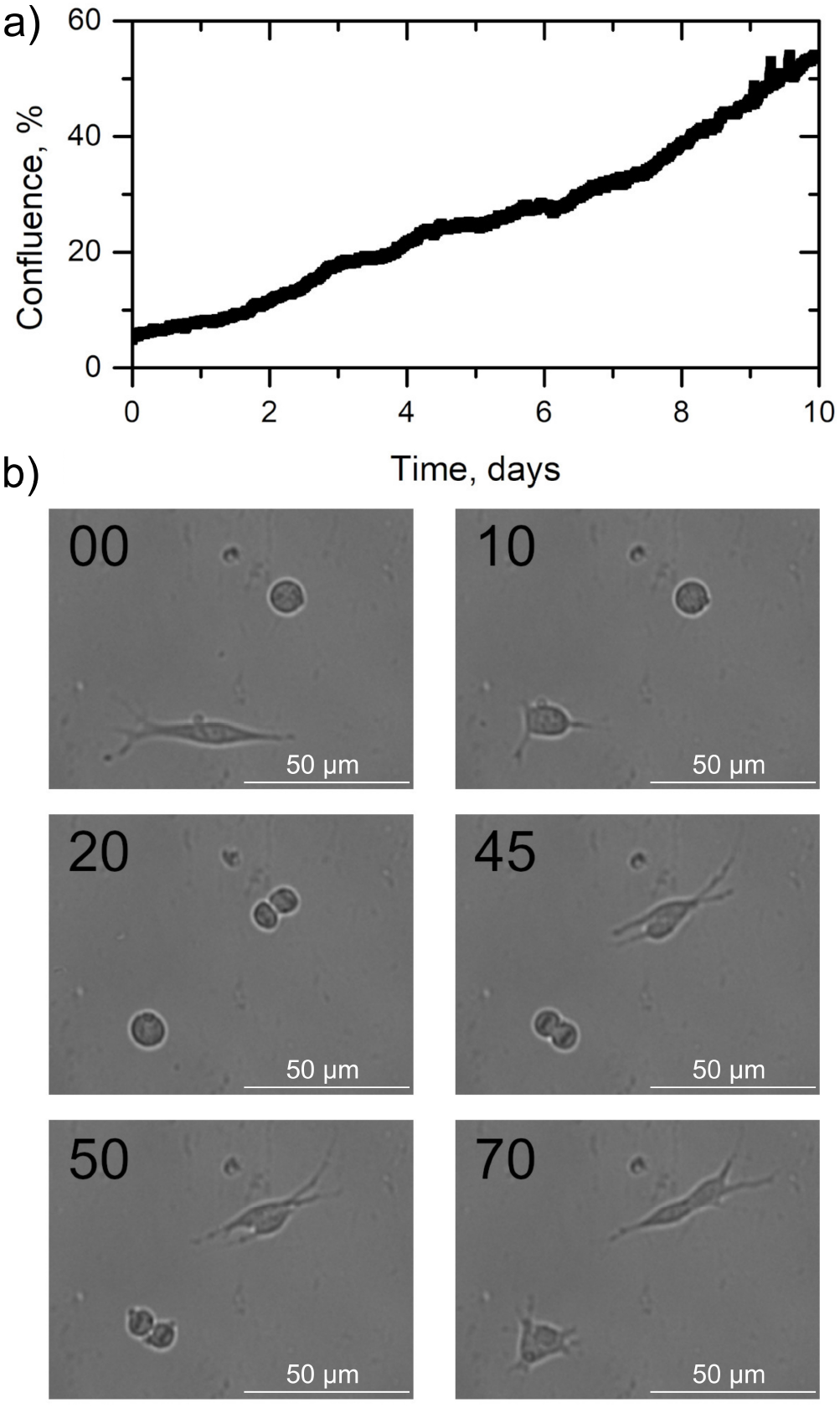
a) The graph illustrates the temporal evolution of surface coverage by SH-SY5Y cells in a sealed polycarbonate chamber. b) The cultivation of SH-SY5Y cells is shown, revealing recognisable morphological transformations - from an elongated surface configuration to a spherical shape - and the distinctive positioning of chromosomal lines within the cell, both before and after division. Each image is temporally annotated in minutes relative to the original photograph, and these individual frames are extracted from a video available in the Zenodo repository.(22)

In addition, the morphological behaviour of SH-SY5Y cells during long-term culture was investigated (see Figure 7b). We observed the morphological transformation of a dividing cell from a stretched surface configuration to a spherical shape. The process of cell division is illustrated in Figure 7b20-b45, showing the positioning of chromosomes and the inclusion of temporal markers to indicate the dynamics of the process. After division, the spherical cells return to their stretched form on the surface. To our knowledge, this morphological feature of SH-SY5Y cell division has not been reported before.

## Conclusions

In this study, we systematically investigated the proliferation of SH-SY5Y cells, HeLaGFP cells, and 16HBE14*σ* cells in polycarbonate chambers with varying surface areas. Our results demonstrated a clear correlation between chamber diameter and cell growth rates, with smaller diameters resulting in slower growth rates for SH-SY5Y and HeLaGFP cells, while 16HBE14*σ* cells proliferated efficiently across different diameters.

The differential responses observed among SH-SY5Y, HeLaGFP, and 16HBE14*σ* cells to varying chamber diameters suggest that cellular mechanisms of adaptation to confined spaces vary significantly between cell types. This observation is novel and suggests potential pathways or regulatory mechanisms that enable certain cells to thrive in confined environments. Understanding these mechanisms could lead to new strategies for controlling cell growth in microfluidic devices and other applications.

The study also showed that SH-SY5Y cells have low mitochondrial function, as evidenced by their reduced oxygen consumption rate. This unique respiratory property makes them ideal for long-term studies in sealed chambers. Our results confirm the practicality of growing these cells in large, sealed chambers (minimum diameter: 10 mm), which simplifies experimental setups and increases the practicality of the microfluidic platform (38).

Additionally, we observed a morphological change in the shape of SH-SY5Y cells during cell division, from stretched to spherical. After division, the cells reverted to their original shape.

Together, these findings provide guidelines for improving experimental design and execution when using SH-SY5Y cells as an in vitro model. By optimising chamber dimensions and exploiting the respiratory properties of the cells, researchers can develop more effective microfluidic platforms for studying neurodegenerative diseases. These advances have the potential to provide better in vitro models for drug screening, toxicity testing and understanding the underlying mechanisms of neurodegeneration, ultimately contributing to the development of novel therapeutic strategies.

## AUTHOR CONTRIBUTIONS

EK: conceptualization, formal analysis, data curation, investigation, methodology, writing – original draft; AL: formal analysis, data curation, investigation, methodology; DW: formal analysis, data curation, investigation, methodology; WS: formal analysis, data curation; JA: formal analysis, data curation; PZ: conceptualization, writing – original draft, and writing – review & editing; SJ: conceptualization, funding acquisition, investigation, methodology, project administration, resources, supervision, writing – original draft, and writing – review & editing.

## COMPETING FINANCIAL INTERESTS

There are no conflicts to declare.

## DATA AVAILABILITY STATEMENT

Two Zenodo repositories(21, 22) contain the original files used to prepare the figures and formulate the conclusions.

## ACKNOWLEDGEMENTS

This research was supported by funding provided by the Program SONATA BIS 9 of the National Science Centre, Poland, Grant No. 2019/34/E/ST4/00281.

## Notes

https://zenodo.org/records/10401181

https://zenodo.org/records/10418022

### Competing Interest Statement

The authors have declared no competing interest.

### Summary of Updates

Figures have been revised in the current version of the manuscript - statistics have been added. The content of the manuscript has been revised according to the reviews received.

